# EEG Correlates of Cognitive Dynamics in Task Resumption after Interruptions: The Impact of Available Time and Flexibility

**DOI:** 10.1101/2024.08.26.609362

**Authors:** Soner Ülkü, Stephan Getzmann, Edmund Wascher, Daniel Schneider

## Abstract

Interruptions are a common aspect of everyday working life, negatively affecting both task performance and long-term psychological well-being. However, research suggests that the effects of interruptions can be mitigated in several ways, such as the opportunity to anticipate the interruptions and preparation time. Here, we used a retrospective visual working memory task to investigate the effects of duration and flexible resumption after interruptions, with 28 participants (18-30 years old) attending the experiment. For this main task participants were required to remember the orientations of a set of colored bars and retrieve one at the end of the trial in response to a retro-cue. This task was sometimes interrupted with an arithmetic task that was presented before the retro-cue. The period after the interruptions and the retro-cue was either short (no additional time), long (additional 1000 ms), or self-determined. Interruptions affected the main task performance irrespective of duration condition, while response times were shorter with the flexible condition. EEG analysis showed that having more time before resuming the interrupted task enabled stronger beta suppression which in turn modulated task performance, helping participants to safely disengage from the interrupting task, and refocus their attention back more efficiently. Further, flexibility in the timing of resumption provided additional benefits as seen in stronger oscillatory alpha and beta suppression to the retro-cue, also being related to better task performance. These results demonstrate the important role of resumption time and individual flexibility in dealing with interruptions.

## Introduction

Being interrupted is not just a part of our daily lives but also of working life in many professions. Studies have shown that interruptions occur multiple times during working hours, and especially with the rising use of technology, they are becoming more and more an integral part of any working environment (Keller et al., 2020; Kiesel et al., 2022; Leroy et al., 2020; Leroy & Glomb, 2018; Puranik et al., 2020). On the level of task performance, they have been linked to decline in the measures of performance such as speed and accuracy of the interrupted task (Arnau et al., 2019; Bae & Luck, 2019; Kiesel et al., 2022; Zickerick, Kobald, et al., 2021; Zickerick, Rösner, et al., 2021). On a more general level, they can also cause anxiety, stress, and even long-term effects on psychological well-being of the workers (Couffe & Michael, 2017; Keller et al., 2020; Kiesel et al., 2022; Leroy et al., 2020). So, it is no surprise that there is an ever-present interest in task interruptions in human factors and basic research literature, with a focus on the underlying cognitive mechanisms and strategies that may reduce the negative consequences arising from task interruptions.

The general definition of an interruption includes (a) a main task a person is attending to, (b) an interrupting task that draws attention and resources away from the main task before it is completed, and (c) the goal to resume the main task after the interrupting task is completed (Couffe & Michael, 2017; Puranik et al., 2020; Zickerick, Kobald, et al., 2021). This requires the flexible use of working memory and attentional control to handle the interrupting task while maintaining the state and the goals of the main task, shift the focus of attention from one task to another, and reactivate the relevant information and task set when the interrupting task is completed. These processes as well as the allocation of mental resources on the interrupting task at the expense of the main task may explain the decline in main task performance (Arnau et al., 2019; Bae & Luck, 2019; Hakim et al., 2020; Lin et al., 2021).

One way to probe this shifting of the focus of attention in working memory is the use of retrospective cuing (retro-cue) paradigms. Such an experimental design would see participants store task-relevant information in working memory before directing attention to a specific subset of information with the presentation of informative cues (Griffin & Nobre, 2003; Schneider et al., 2017; Souza & Oberauer, 2016). It is known that this retro-cue paradigm provides a general benefit for visual working memory performance, while this focusing of attention in working memory cannot compensate for deficits arising from interruptions. In other words, such a retro-cue paradigm is well suited to study the temporal course of the shift of attention without any confounding effect, and how attentional control processes on the level of working memory are affected by prior interruptions (Gilchrist et al., 2016; Rösner, Sabo, et al., 2022; Schneider et al., 2017; Zickerick, Rösner, et al., 2021).

The impact of interruptions on working memory may vary depending on multiple task characteristics. Firstly, tasks that are less cognitively demanding or cause less competition for working memory capacity are less affected by interruptions compared to tasks that are more complex or induce sudden shifts of attention (Altmann et al., 2014; Cades et al., 2007; Monk et al., 2008; Zickerick, Rösner, et al., 2021). Furthermore, certain task parameters are found to help mitigate the negative effects of interruptions on working memory. Earlier studies on multitasking showed that the time it takes to switch between tasks decreased significantly if participants are given a preparatory period before starting a new task. Likewise, negative consequences of interruptions on task performance can be mitigated if participants can anticipate an upcoming interruption (Kiesel et al., 2010; Labonté et al., 2019; Meiran et al., 2000; Trafton et al., 2003; Ülkü et al., 2024). It is also known that the general predictability of temporal structures of the upcoming tasks does not just boost the main task performance, but also the performance in the interrupting task (Gresch et al., 2021, 2022). Further, studies using the retro-cue paradigm indicated that having more time before or after the presentation of a retro-cue can increase the main task performance, probably by giving more time to focus attention on relevant information and plan upcoming motor responses (Liu et al., 2023; Schneider et al., 2016; Shin et al., 2017; Wang et al., 2018). An open question in current literature, however, is the comparison of these general timing effects with a self-determined task resumption. In real world scenarios, task switching and interruptions occur rarely with rigid timing, instead there is more autonomy of the individual on how and when to proceed with an interrupted task.

To examine if and how self-pacing of task resumption might help counteract the interruption-based deficits, we implemented an experimental design which enabled participants to actively control when to resume with the main task after being interrupted. The main task consisted of a visual working memory task in which participants had to maintain the orientation of two out of four different bars. After the maintenance period, a retro-cue indicated which of the two bars’ orientation had to be reported. During half of the trials, participants were interrupted with a visual arithmetic task before they could report the cued item. To probe the effects of timing and how self-paced task resumption might help deal with interruptions, we established a fixed and a flexible timing condition. In fixed trials, the onset of the main task, the onset of the interrupting task, and the resumption of the main task was fixed and predefined. In contrast, in flexible interruption trials, participants were able to choose when to resume the main task. In addition, to investigate the role of the time provided to resume the main task, the fixed timing trials (both interruption and no-interruption) were further divided into short and long duration. Specifically, the long trials had an additional 1000 ms after the offset of the interrupting task.

In order to investigate the underlying neuro-cognitive mechanisms when dealing with interruptions, and to explore the effects of self-paced resumption and additional duration after an interruption, we employed electrophysiological measurements and time-frequency decompositions. We concentrated on mid-frontal theta oscillations (4 – 7 Hz) which can be used as a measure of attentional control processes and mental resource allocation (Cavanagh & Frank, 2014; Cavanagh & Shackman, 2015; Riddle et al., 2020; Senoussi et al., 2022; Zickerick, Rösner, et al., 2021), and posterior alpha oscillations (8 – 14 Hz), which are known to be affected by the manipulation of working memory content and the focusing of attention in working memory (De Vries et al., 2020; Myers et al., 2015; Schneider et al., 2022; Zickerick, Rösner, et al., 2021).

We expected to see reduced performance of the main task when participants were interrupted, in line with previous studies (Arnau et al., 2019; Bae & Luck, 2019; Rösner, Zickerick, et al., 2022; Zickerick, Kobald, et al., 2021). However, these effects should be reduced for the trials where participants were able to resume the main flexibly (i.e., in a self-paced way). We further hypothesized that having more time would aid the primary task performance in contrast to being forced to immediately resume the main task following an interruption. For the neuro-cognitive measures, we expected to replicate the previous findings of lower evoked theta oscillations and weaker alpha suppression after the retro-cue in interruption, than non-interruption, trials (Rösner, Zickerick, et al., 2022; Ülkü et al., 2024; Zickerick, Rösner, et al., 2021). Moreover, these effects should be reduced in the flexible trials compared to the fixed resumption trials, which would indicate that the self-determined resumption of the primary task after an interruption would facilitate attentional control processes.

## Methods

### Participants

There was a total of 31 participants attending the experiment (age: 18-30). Two of them did not complete the experiment due to personal reasons, and data from one participant had to be discarded due to technical problems. Further, two participants were rejected due to not performing above the chance level on the interrupting task, and one participant due to high amount of rejected trials from EEG processing (see below for more details), leaving 25 participants for analysis (*M_age_±SD* = 24.16±3.35 years, 11 females, 14 males). The participants were right-handed (according to an adapted version of Edinburgh Handedness Inventory), and they reported no known neurological or psychiatric disorders. Further, they were tested for normal color vision using Ishihara Test for Color Blindness. The compensation was 12 € per hour or an equivalent amount of course credits, and each participant gave written informed consent. The study was approved by the ethics committee of the Leibniz Research Centre for Working Environment and Human Factors and was in accordance with the Declaration of Helsinki.

### Experimental Procedure

Participants completed a set of neuropsychological tests (TMT Part A & B, Color-Word-Test, and Number Repetition Forwards & Backwards) before the experiment was started. They were asked to sit in front of a 21-inch CRT monitor at a viewing distance of approximately 110 cm (refresh rate: 100 Hz, resolution: 2048*1536). The programming of the paradigm was done in Lazarus IDE (Free Pascal), and the stimuli were presented via ViSaGe MKII Stimulus Generator (Cambridge Research Systems, Rochester, UK).

The main task was a visual working memory task, in which the orientations of colored bars had to be memorized and later reported. The interrupting task was a two-alternative forced-choice task, in which the participants had to decide if the presented summation was correct or not. Trials were organized into separate “fixed timing” and “flexible timing” blocks, and each block began with the announcement of the condition.

During fixed timing blocks, participants had to complete four different types of trials. They were either interrupted or not and the duration between the presentation of the memory array and the retrospective cue (retro-cue) was either short or long. This also means that the resumption phase after offset of the interruption task and before presenting the retro-cues was either short (500 ms) or long (1500 ms). In all trials, participants were first presented with a memory array on a dark background (color in CIE1931 space: 0.287, 0.312, 15), consisting of two randomly oriented blue bars (CIE1931: .195, .233, 42; size: 1° * 0.1°) presented laterally and placed at the center of their respective quadrants of the screen, and two randomly oriented orange bars on the opposite side (CIE1931: .484, .451, 42; size: 1° * 0.1°). The placement of the blue vs. orange bars on the laterals was done randomly on each trial. Each participant was given one specific color to focus on for the whole duration of the experiment and was asked to remember their orientation. Target color was counterbalanced across participants. These four items were presented for a duration of 200 milliseconds and were replaced with a centrally presented fixation cross (CIE1931: .287, .312, 40; length of bars: 0.95°). In no-interruption trials, this fixation cross was presented for a total of 3800 ms in the short duration condition and for 4800 ms in the long duration condition. It was then replaced by a retro-cue that was presented centrally for 200 ms (height: 1.23°), which indicated if the upper or lower bar from memory array subset would be relevant for the orientation report. In interruption trials, this initial fixation cross was replaced by a centrally presented equation for 1800 ms (digit height: 0.95°), with two single digit numbers summing up to a two-digit number. During this presentation participants had to report if the summation was correct or not by pressing the left or right computer mouse button. This response-button mapping was counterbalanced across participants. After this interrupting task, participants were presented with the fixation cross for 500 ms in the short duration condition and for 1500 ms in the long duration condition. This retro-cue was followed by the presentation of an additional fixation cross for 800 ms, which was replaced by a memory probe (CIE1931: .287, .312, 0; size: 1° * 0.1°) in random orientation. Participants then had to use the computer mouse to adjust the orientation of this memory probe by moving the mouse on a horizontal axis, so the orientation would match the orientation they had to report. To finalize their answer, participants were asked to click on the left mouse button. This final stimulus presentation was replaced with a fixation cross for the duration of the inter-trial interval (500-1000 ms) after participants responded, or if no response was given, after a maximum of 4000 ms.

During the “flexible timing” blocks, the no-interruption trials were structured the same way, and only differed in their duration (short vs long). However, in case of an interruption, participants were able to flexibly choose the time until they proceeded with the main task. Thus, following their response to the interrupting task, participants had no fixed time limit, but could indicate by a button press when they wanted to resume the main task. After this second response they were presented with a fixation cross for 500 ms, which was followed by the usual structure of the trials in the order of retro-cue presentation, additional fixation, and memory probe presentation. In total there were 360 trials with interruptions (120 short duration, 120 long duration, 120 flexible timing trials), and 360 trials without interruptions (180 short duration, 180 long duration trials), which were distributed across 9 blocks (3 flexible timing, 6 fixed timing).

### Behavioral Data

Only trials in which participants responded to the primary task (and to the interrupting task if there was one), and in which response times to the interrupting task were above 150 ms (since the others were considered as premature responses) were included in the further analyses. Additionally, to make them comparable to the other conditions, flexible resumption trials were only accepted if the responses to the interrupting tasks were given within 1800 ms. Further, to match the EEG and behavioral parameters, the trials used for behavioral analysis were only considered if they were also included as EEG trials. Two performance parameters were derived from each task: for the main task, the angular error (i.e., the absolute degree difference between the actual target bar’s orientation and the reported orientation) and the response onset time (i.e., the time between the presentation of the memory probes and the moment when the participants started to move their mouse). For the interrupting task, the response times and task accuracy were assessed (i.e., whether the response to the arithmetic task was correct). This last measure was calculated by dividing the total number of correct responses by the respective number of trials of a given condition. Since during the flexible timing interruptions, participants were theoretically able to give “late” responses to the interrupting task, these were treated as misses and not included for the statistics. This was done to match the interrupting task itself and the responses it induced across conditions, and not use the trials in which the participant might have used the additional time in flexible condition to further focus on the interrupting task. Since there was one factorial combination missing (i.e., the flexible no-interruption condition), two different ANOVAs were performed: First, a repeated-measure ANOVA was run on the behavioral measures, using duration (short vs long), and interruption (no-interruption vs interruption) as independent variables, leaving out the flexible resumption trials for this analysis. Secondly, an additional one-way ANOVA was run only for the interruption trials using duration (short vs long vs flexible) as a repeated-measures factor. Bonferroni-Holm correction for multiple comparisons was used for further post-hoc t-tests.

### Electrophysiological Data

The EEG was recorded using a 64-channel passive electrode cap (Easycap 64 Ch-Braincap, Easycap GmbH, Herrsching, Germany; extended 10/20 scalp configuration; sampling rate: 1 kHz). The ground channel was AFz and the reference channel was FCz. The signals were amplified via NeurOne Tesla TMS Amplifier (Bittium Bio-Signals Ltd, Kuopio, Finland). This recorded data was then processed with EEGLAB v2022.1, running on MATLAB v9.14.0 (R2023a, Mathworks, Natick, USA).

First, the data was band-pass filtered with cut-off frequencies of 1 and 45 Hz (pop_eegfiltnew, FIR, Hamming, high-pass filter order = 3300, low-pass filter order = 330). Then channels were rejected on the premise of flat line (clean_flatlines) or extreme noise (pop_rejchan, SD = 5) using kurtosis as a criterion. In the end 4.32 channels were rejected on average (SD = 2.15). Data was then re-referenced to average, and the missing channels were interpolated (pop_interp, spherical). Additionally, Independent Component Analysis was used to clear out artefacts arising from eye blinks and external noise. A copy of the continuous data was epoched into 9 second trials relative to the memory array onset, starting 1000 ms before and ending 8000 ms after. Trials were rejected and cleared via an automated trial rejection procedure (pop_autorej, threshold: 1000 µV, prob. threshold: 5 SD, max. rejection of trials per iter.: 5%). The cleared data was downsampled to 250 Hz, and an ICA algorithm was run on this dataset (infomax, sub-Gaussian sources, PCA). Another algorithm was run on these components to classify and organize them, removing components which had a probability higher than 50% to reflect eye movements, eye blinks, line noise, or channel noise. In the end, 4.72 components were rejected on average (SD = 2.16). The resulting component weights and labels were transferred to the original dataset which was epoched like the ICA data. As a final step of cleaning, trials were rejected if they had large fluctuations (pop_eegthresh, th_low = -150 µV, th_high = 150 µV), resulting in an average of 40.52 trials rejected from the data (SD = 43.56).

### Oscillatory Power

To investigate the changes in oscillatory power, frequencies in logarithmic steps in the range of 4 – 30 Hz were extracted using wavelet convolution per channel with a Full-Width-at-Half-Maximum range of 750 to 100 ms. The baseline was selected to be from 500 ms before to 200 ms before the memory array onset, providing a condition-general oscillatory baseline. The time-frequency decomposition of the trials resulted in event related spectral perturbations (ERSPs) from 700 ms before to 7660 ms after the memory array onset. These decompositions were then down sampled to 25 Hz to ease the calculation and data storage. Since the “flexible” condition induced time variability following interruption offsets, an additional time-frequency decomposition was run on epochs time-locked to the retro-cue onset, aligning all conditions temporally. This secondary time-frequency decomposition resulted in ERSPs from 1700 ms before to 1680 ms after retro-cue onset, which we used for our statistical analysis. To probe the effects of interruption, duration, and self-determination of primary task resumption, we made use of cluster-based permutation tests using FieldTrip, for a given time window around the retro-cue where the conditions shared the general structure of the trial (500 ms before the retro-cue to 1000 ms after the probe for fixed trials; from the onset of the retro-cue to 1000 ms after for the comparison of flexible and fixed interruption trials). The following parameters were used for cluster-based permutation statistics in FieldTrip, *estimation method*: Monte-Carlo, *correction method*: cluster, *alpha level for a cluster to be significant*: 0.05, *parameter to collect from the clusters*: maximum of the t-statistic sum, *minimum number of channels for a cluster to be defined*: 2, *alpha to control for the false alarm rate*: 0.05, *number of randomizations*: 1000. The function ft_prepare_neighbours was used to define neighbor relations of channels. The clusters were selected as significant if their p-values were lower than 0.05, which was calculated by finding the percentile where the t-sum of the said cluster was positioned on the aggregated maximum of t-sum values over permutations. Since each significant cluster resulted in a large amount of significant time–frequency points for several channels, we provide a metric indicating how much each channel contributed to the overall cluster. This metric is simply the ratio of the significant datapoints per channel, normalized by the number of datapoints from the channel contributing the most to the cluster (see figures 4 to 6).

### Software Packages

NumPy (v. 1.26.4) and Pandas (v. 2.2.1) packages were used for handling the data and descriptive statistics, Pingouin (v. 0.5.4) for ANOVAs and post-hoc t-tests, and Matplotlib (v. 3.8.0) and seaborn (v. 0.13.2) for data visualization (Harris et al., 2020; Hunter, 2007; McKinney, n.d.; Vallat, 2018; Waskom, 2021).

## Results

### Behavioral Data

We focused on two performance parameters of the main task, (a) the angular error (i.e., the average difference between the actual target bar’s orientation and the participant’s response in degrees), and (b) the response onset (i.e., the time when the participants started moving their mouse to execute their response) as a correlate of the actual response times in a continuous report working memory task (Figure 2).

**Figure 1.**
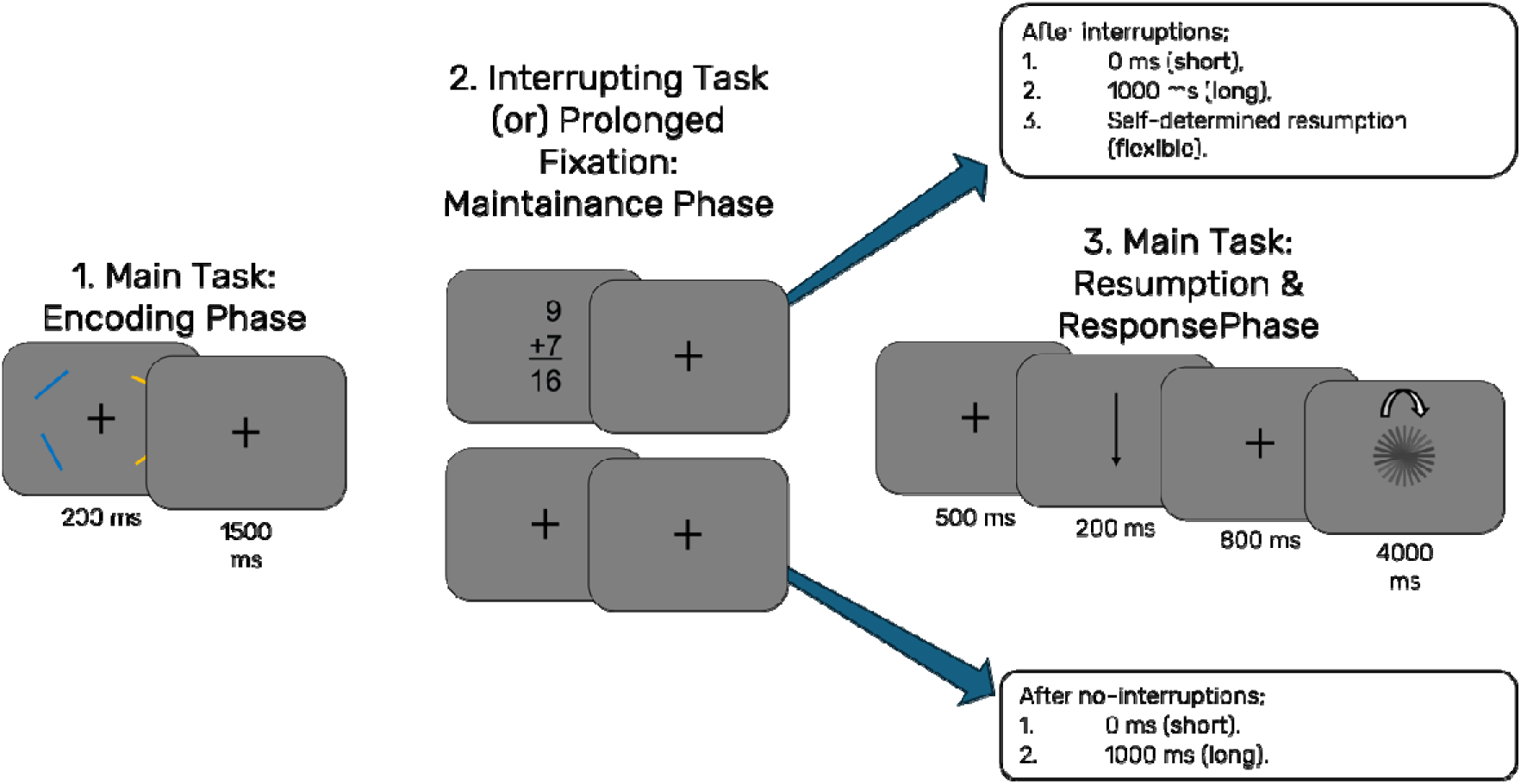
Experimental design. All participants were assigned a target color at the beginning of the experiment and were instructed to store only the orientation of the bars presented in this color (orange vs. blue). After the participant was presented with the memory array, there was a delay of 1500 ms with a fixation cross on the screen before they were presented with the interrupting task for 1800 ms or an additional fixation cross for the same duration. After this, depending on the trial type, either (1.) the task continued to the resumption phase, (2.) the participants had a further 1000 ms of fixation period, or (3.) they had unlimited time to self-report when they want to resume the task. The resumption phase included a fixation period of 500 ms, a retro-cue that was presented for 200 ms, an additional fixation period of 800 ms, and a presentation of a memory probe that was dismissed either via participant response or after a duration of 4000 ms.

**Figure 2.**
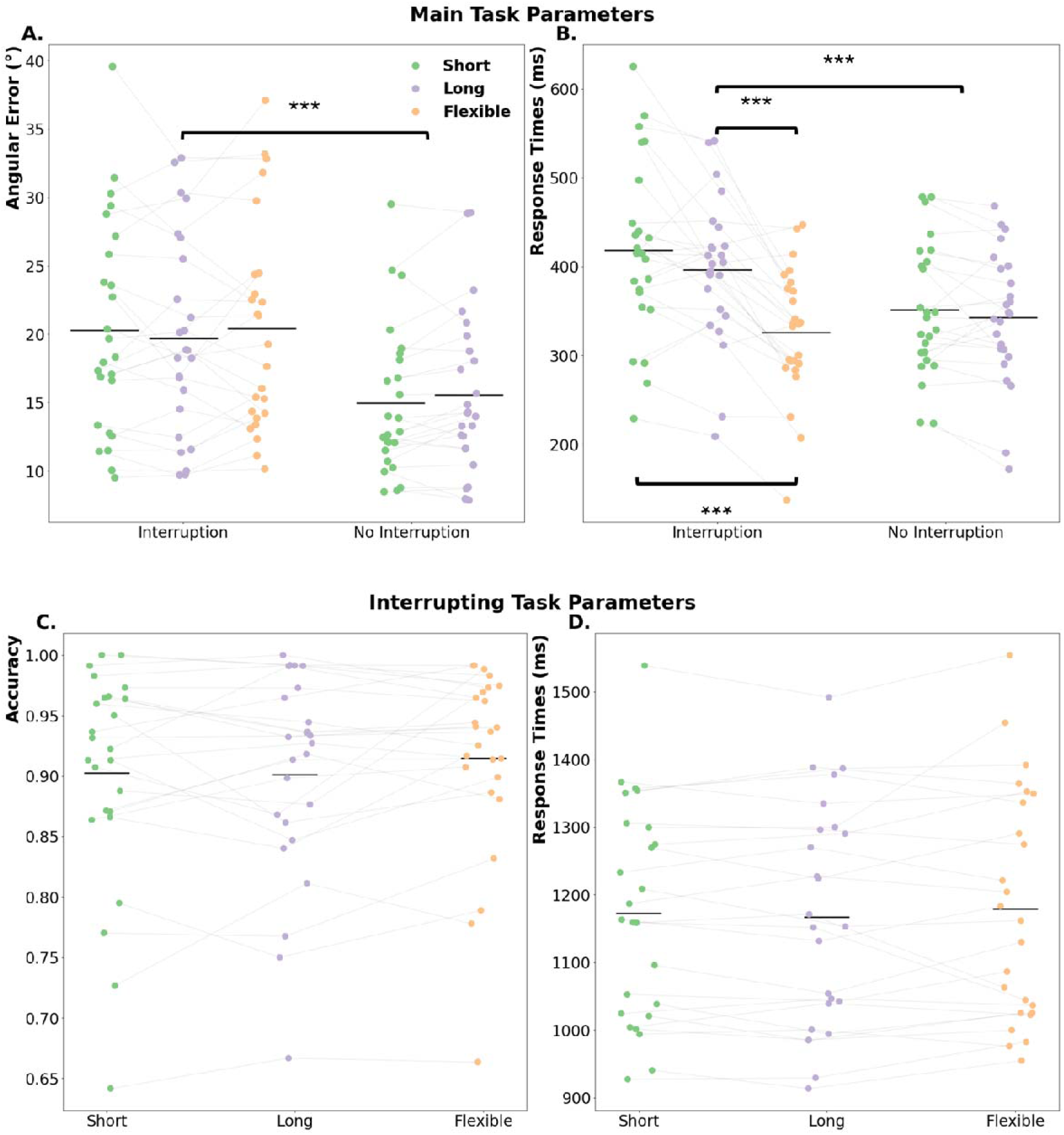
Behavioral parameters from the primary task. (A) Angular error, (B) precision and (C) response onset times are shown separately for interruption and no-interruption trials, and for each duration condition. The colored dots represent the individual values, and the horizontal lines indicate mean values for the given parameter. The asterisks represent the significant differences between experimental conditions (p<.001 ***, p<.01 **, p<.05 *).

**Figure 3.**
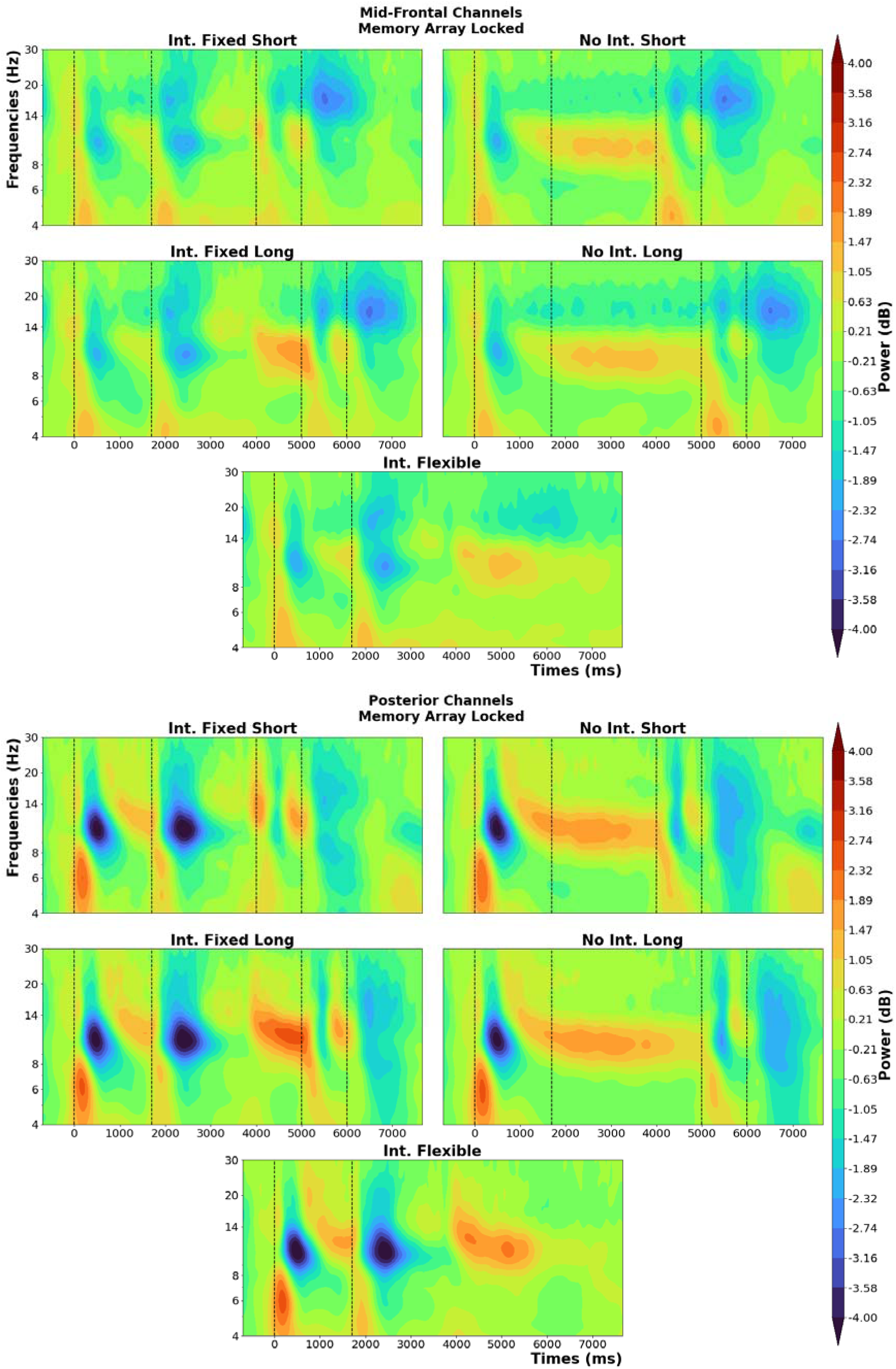
Time-frequency plots per condition, with the baseline corrected oscillatory power measures. The upper four subplots are from a cluster of mid-frontal channels (Fz, F1, F2, FC1, FC2) and the lower five subplots are from a posterior channel cluster (PO3, POz, PO4, O1, Oz, O2). The events (onsets of memory array, interruption, retro-cue, and probe) are marked by vertical dotted lines, and their timing changes depending on condition.

As expected, interruptions reduced main task performance, which was confirmed with a main effect of interruptions on both accuracy, *F*(1,24)□=□55.63, *p*□<□.001, η*_p_^2^*□=□.7, and on response times, *F*(1,24)□=□20.99, *p*□<□.001, η*_p_^2^*□=□.47 (see Figure 2A, Interruption: *M±SD* = 20.15°±7.48°, No-Interruption: *M±SD* = 15.26°±5.59°; Figure 2B, Interruption: *M±SD* = 380.43±91.64 ms, No-Interruption: *M±SD* = 346.99±72.89 ms). Further, there was a performance benefit stemming from flexibility and longer resumption periods regarding response times that was confirmed with a main effect of duration for fixed trials, *F*(1,24)□=□4.92, *p*□=□.04, η*_p_^2^*□=□.17, and for interruptions only trials, *F*(2,48)□=□32.43, *p*□<□.001, η*_p_^2^*□=□.57 on response times (see Figure 2B; Short: *M±SD* = 384.77±92.03 ms, Long: *M±SD* = 369.74±80.61 ms, Flexible: *M±SD* = 326.25±71.94 ms).

Running post-hoc t-tests revealed that the condition with interruptions followed by a flexible resumption phase had significantly faster responses to the main task, compared to both short (*t*(24) = 6.55, *p_corr_* < .001, *cohen’s d* = 1.08, 95% CI [63.2, 121.33]) and long duration conditions (*t*(24) = 6.80, *p_corr_* < .001, *cohen’s d* = 0.93, 95% CI [48.96, 91.65]). The comparison between long and short resumption phases showed no significant difference in response times (*t*(24) = 1.97, *p_corr_* = .06, *cohen’s d* = 0.25, 95% CI [-1.08, 45.0]).

We additionally wanted to see if these fast responses in the flexible resumption condition were merely stemming from using the time before the retro-cue to prepare the upcoming motor response. To this end we ran correlations between these resumption lags (i.e., the time participants took before resuming the main task) and the working memory task accuracy and response times on a trial-by-trial basis for each participant. We then tested the Z-transformed correlation coefficients against zero. This showed that participants had a trend – albeit insignificant – for higher accuracy on the working memory task if they took less time to resume this task after an interruption (*t*(24) = 1.96, *p* = .06, *cohen’s d* = 0.39, 95% CI [0., 0.09]). Furthermore, there was no reliable single-trial correlation between resumption phase and working memory task response times (*t*(24) = 0.77, *p* = .45, *cohen’s d* = 0.15, 95% CI [-0.03, 0.07]).

The performance in the interrupting task did not show an effect of duration, neither for accuracy with *F*(2,48)□=□1.69, *p*□=□.2, η*_p_^2^*□=□.07 (see Figure 2C, Short: *M±SD* = 90±9%; Long: *M±SD* = 90±8%; Flexible: *M±SD* = 91±8%), nor for response times *F*(2,48)□=□0.77, *p*□=□.47, η*_p_^2^*□=□.03 (see Figure 2D , Short: *M±SD* = 1172.87±160.13 ms; Long: *M±SD* = 1167.27±164.12 ms; Flexible: *M±SD* = 1179.24±170.82 ms).

### Electrophysiological Data

To make it easier to follow the description of the EEG results, we first provide descriptive oscillatory plots that shows neural oscillations time-locked to the memory-array for a group of mid-frontal and posterior channels.

The cluster-based permutation was run similarly to the way we implemented ANOVA for the behavioral analyses. For the fixed trials, these stats only included the time range 500 ms before the retro-cue up until the presentation of the memory probe to exclude any oscillatory activity stemming from motor responses to the interrupting task or the main task. The initial contrast was between interruption and no-interruption conditions, including only fixed duration trials (see Figure 4, top row). The first cluster (*t-*sum = 212438.41, critical *t* = 16982.25, *p* = .001) seems to be around the alpha and beta frequency bands, with a posterior topography and a timing that starts before the onset of the retro-cue and continues until the presentation of the memory probe. It indicated that participants showed stronger alpha/beta suppression around and after the retro-cue when they were not interrupted. The second cluster (*t-*sum = -35430.08, critical *t* = -16008.13, *p* = .01) had a narrower distribution over the theta frequency band and started right before the onset of the retro-cue and ceased before the presentation of the memory probe. The effect was strongest around left mid-frontal channels and indicated that participants exhibited higher oscillatory power in the theta frequency range to the retro-cue when they had not been interrupted.

**Figure 4.**
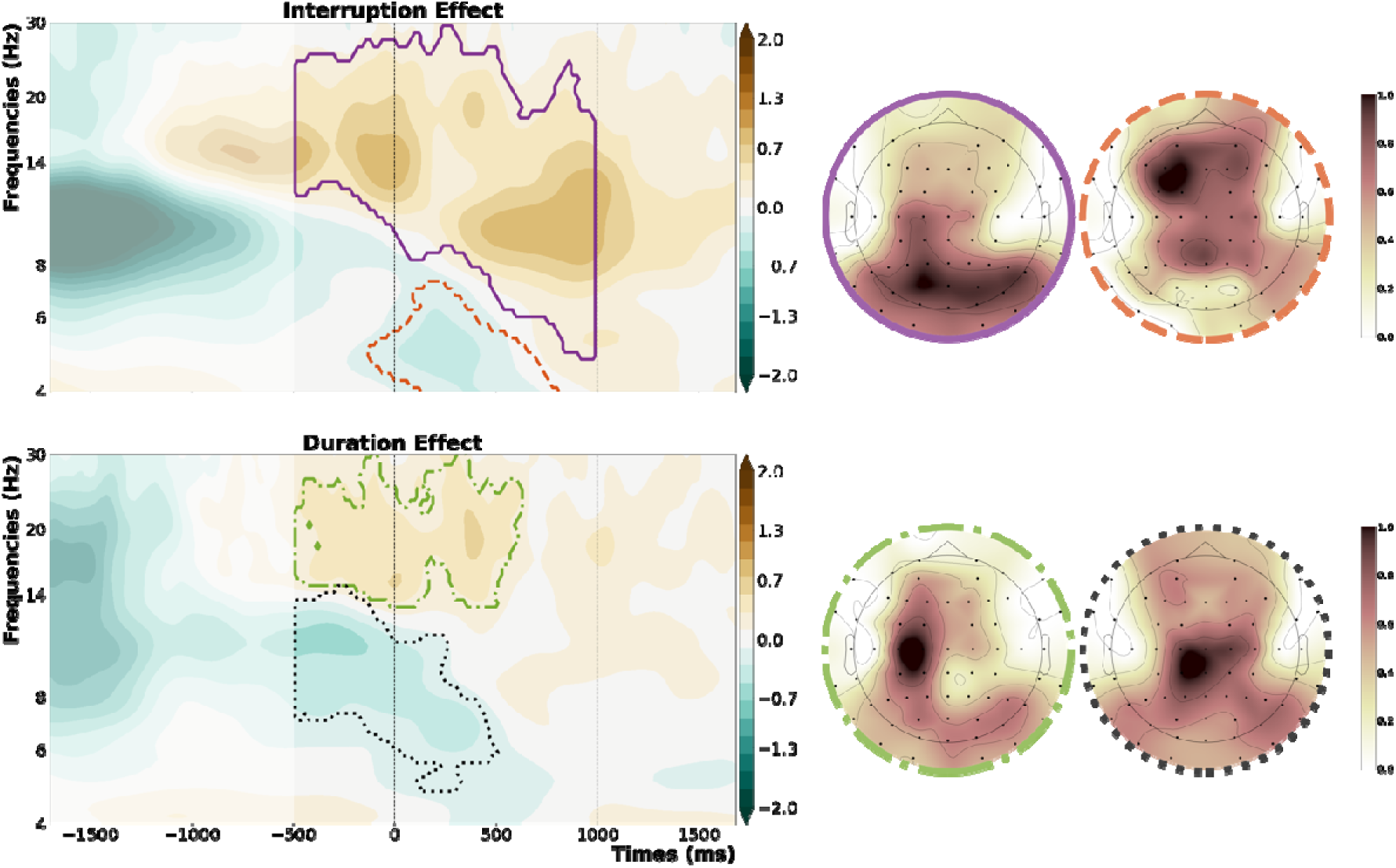
Significant clusters showing the main effects of interruption and duration, plotted separately. First row shows the interruption effect with the topographies marking the specific cluster’s distribution over the scalp. Second row on the bottom shows the duration effect, with the vertical dashed lines representing the onset of the retro-cue (0 ms) and the onset of the memory probe (1000 ms).

Following the interruption effects, we compared long duration trials with short duration trials (see Figure 4, bottom row). Here, the first cluster (*t-*sum = 67695.63, critical *t* = 13223.64, *p* = .001) was centered around beta frequency band and its topography was localized at the left-hemispheric motor cortex, starting already before the onset of the retro-cue and indicating stronger beta suppression around the retro-cue for trials that had a longer maintenance phase. The second cluster (*t-*sum = -68546.61, critical *t* = -13685.99, *p* = .003) was localized around the alpha frequency band and started already before the onset of the retro-cue. It had a narrow distribution over left parietal channels, indicating higher oscillatory power before the retro-cue for trials with a longer duration.

To see if there was any interaction between the effects of interruptions and duration, we subtracted the oscillatory power of longer trials from shorter trials, time-locked to the retro-cue, and compared this between interruption and no-interruption conditions (see Figure 5). This comparison revealed three significant clusters, with three distinct topographies. The first cluster (*t-*sum = 13389.01, critical *t* = 11765.64, *p* = .05) appeared across the oscillatory beta band, starting before the presentation of the retro-cue. It indicated a stronger duration effect on beta oscillations for interruption trials, with a topography over left motor cortex. The second cluster (*t-*sum = 12177.88, critical *t* = 11765.64, *p* = .05) showed a stronger duration effect on theta oscillations for interruption trials than no-interruption trials, with a left mid-frontal topography. The third cluster (*t-*sum = -117116.93, critical *t* = -13971.53, *p* = .001) appeared across the alpha band and showed a stronger duration effect for no-interruption trials, with the strongest effect at right parietal/ posterior channels.

**Figure 5.**
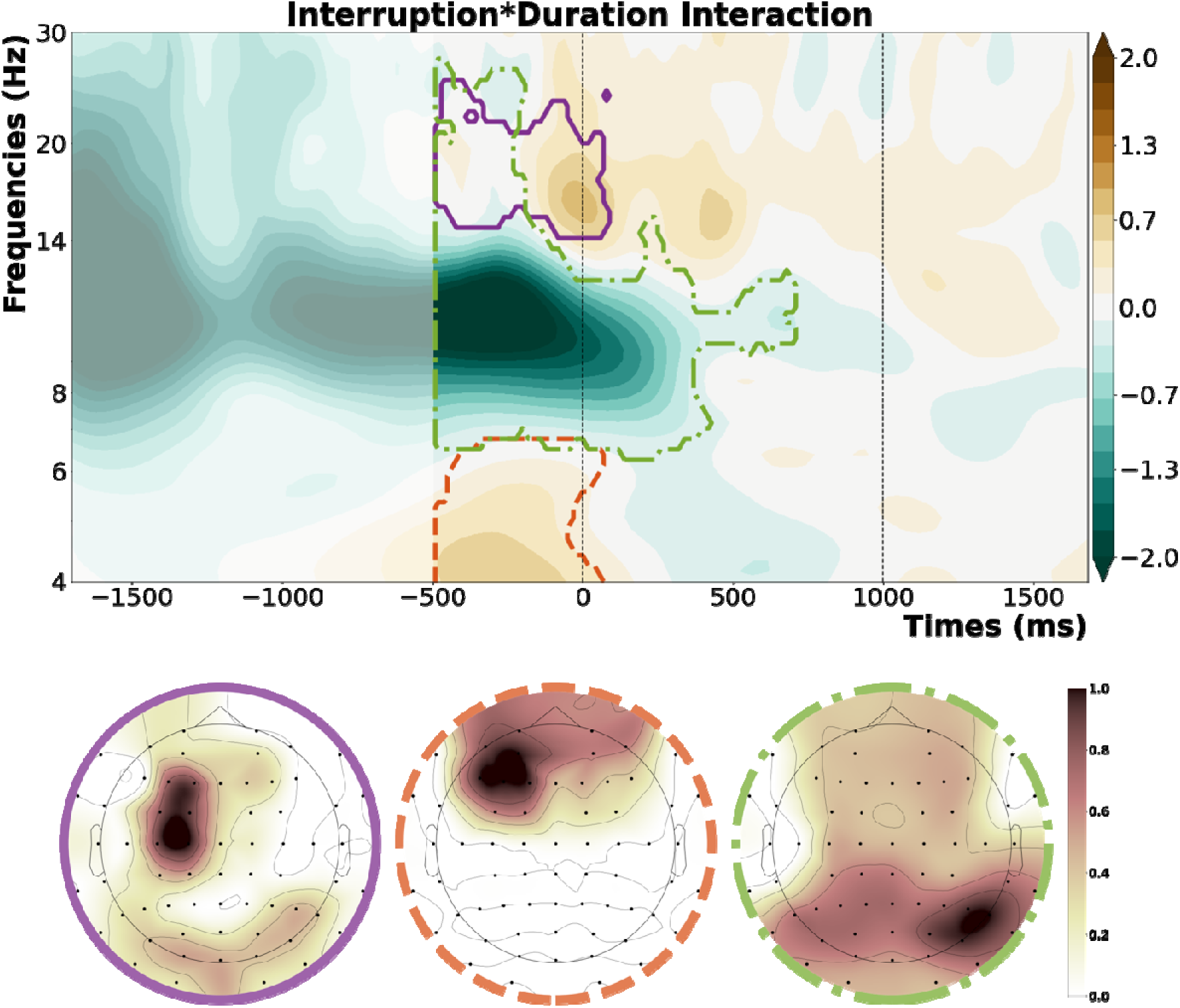
Significant clusters showing an interaction between the effects of interruption and duration, the first row shows the contour-plot for the three clusters, with the vertical dashed lines representing the onset of the retro-cue (0 ms) and the onset of the memory probe (1000 ms). The second row on the bottom shows the topography of each cluster.

**Figure 6.**
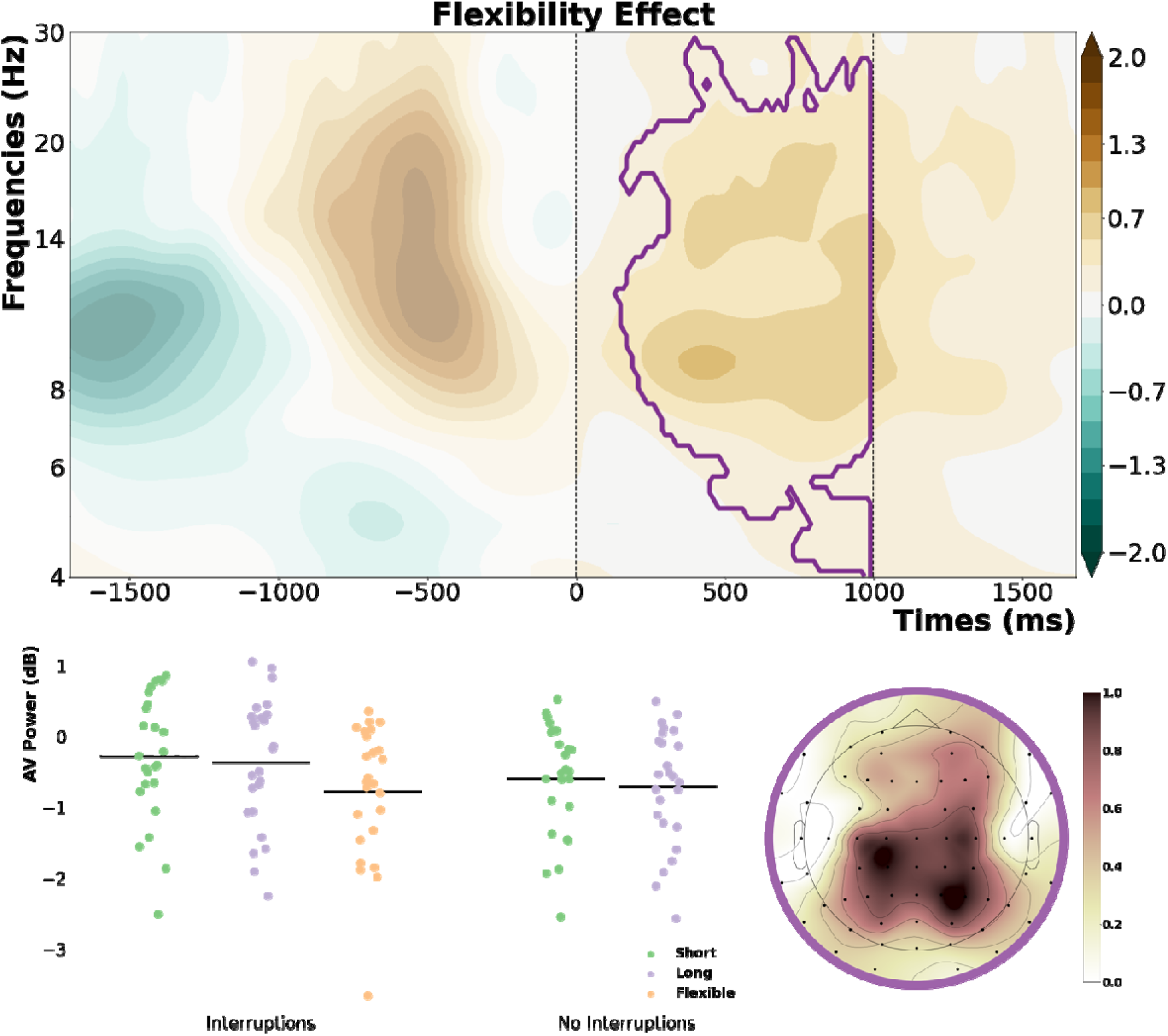
Significant cluster which showed the main effect of flexibility. The contour plot shows the oscillatory power difference between flexible and fixed interruptions, with the vertical dashed lines representing the onset of the retro-cue (0 ms) and the onset of the memory probe (1000 ms). The scatterplots show the individual power values per subject within this cluster for each condition, and the topography shows the distribution of the effect.

To further disentangle these interactions, we compared short duration trials to their longer counterparts separately for interruption and no-interruption conditions, by averaging the oscillatory power within these clusters and running t-tests on the resulting parameters. This revealed that the effect within the beta cluster was mainly driven by the interruption conditions, where we were able to see a significant effect of duration (*t*(24) = 6.30, *p_corr_* < .001, *cohen’s d* = 0.79, 95% CI [0.48, 0.95]), with lower oscillatory power in the interruption condition with longer resumption phase. There was no significant difference within the no-interruption condition (*t*(24) = 1.9, *p_corr_* = .07, *cohen’s d* = 0.12, 95% CI [-0.01, 0.21]). For the theta cluster, the interruption condition showed a significant effect of duration (*t*(24) = 4.36, *p_corr_* < .001, *cohen’s d* = 0.98, 95% CI [0.36, 1.01]), with stronger oscillatory power in the interruption condition with shorter resumption phase. This was again not the case for the no-interruption condition (*t*(24) = -1.69, *p_corr_* = .1, *cohen’s d* = 0.18, 95% CI [-0.27, 0.03]). However, we were able to see that the duration effect within the alpha cluster was significant for both the interruption condition (*t*(24) = -4.81, *p_corr_* < .001, *cohen’s d* = 1.04, 95% CI [-1.22, -0.49]), and no-interruption condition (*t*(24) = 5.36, *p_corr_* < .001, *cohen’s d* = 0.35, 95% CI [0.17, 0.39]), with the sign or the direction of the observed effect changing for the no-interruption condition.

The last step of the cluster-based permutation was a comparison between fixed and flexible interruptions (see Figure 6). Since the resumption was initiated by a motor response 500 ms before retro-cue onset, we only included the time range starting from the onset of the retro-cue until the presentation of the memory probe. This revealed one significant cluster (*t-* sum = 113345.14, critical *t* = 10139.16, *p* = .001), which was centered around the alpha and beta frequency bands and started shortly after the onset of the retro-cue. Its topography had a posterior/parietal distribution and indicated stronger alpha/beta suppression for flexible interruptions. To further disentangle this effect between short, long, and flexible conditions; we implemented post-hoc tests by averaging individual oscillatory power within this cluster for each participant and condition and running multiple comparisons on these values. The post-hoc t-tests revealed that oscillatory power for interruptions with a flexible resumption phase differed significantly from both short (*t*(24) = -4.28, *p_corr_* = .001, *cohen’s d* = 0.54, 95% CI [-0.74, -0.26]), and long duration conditions (*t*(24) = -4.47, *p_corr_*= .001, *cohen’s d* = 0.44, 95% CI [-0.6, -0.22]), whereas short and long duration conditions did not differ from each other (*t*(24) = 1.26, *p_corr_* = .22, *cohen’s d* = 0.09, 95% CI [-0.06, 0.23]). As shown in figure 6, it is also interesting to note that oscillatory power in this time-frequency cluster did not differ between the flexible condition and the no-interruption conditions (*t*(24) = -1.81, *p_corr_*= .17, *cohen’s d* = 0.14, 95% CI [-0.27, 0.02]).

Finally, to see if there was any relation between the EEG parameters and behavioral performance, we made use of the time-frequency clusters from the *interruption * duration* interaction and the flexibility effect and compared the average oscillatory power between good and bad performance by a median split based on the single-trial angular error in the working memory task for each participant. This exploratory analysis only showed significant results for the beta cluster (see figure 5), but only for interruption trials followed by long resumption phases (Good vs Bad: *t*(24) = -3.01, *p_corr_* = .02, *cohen’s d* = 0.28, 95% CI [-0.44, -0.08]). The other conditions on the other hand showed no effect, neither for interruptions followed by short duration resumption phases (Good vs Bad: *t*(24) = -1.48, *p_corr_* = .46, *cohen’s d* = 0.15, 95% CI [-0.34, 0.06]), nor for the no-interruption conditions (short duration: *t*(24) = -1.02, *p_corr_* = .46, *cohen’s d* = 0.09, 95% CI [-0.24, 0.08]; long duration: *t*(24) = -1.31, *p_corr_* = .46, *cohen’s d* = 0.12, 95% CI [-0.28, 0.06]).

When we used the same exploratory approach for the relation between oscillatory power within the flexibility cluster (see figure 6) and behavioral performance, there was a significant relation in the short resumption condition (Good vs Bad: *t*(24) = -3.44, *p_corr_* = .01, *cohen’s d* = 0.28, 95% CI [-0.41, -0.1]) and flexible resumption condition (Good vs Bad: *t*(24) = -2.94, *p_corr_* = .01, *cohen’s d* = 0.21, 95% CI [-0.35, -0.06]). There was only a trend into the same direction for interruptions with longer resumption phases (Good vs Bad: *t*(24) = -1.95, *p_corr_* = .06, *cohen’s d* = 0.14, 95% CI [-0.26, 0.1]).

## Discussion

Here we investigated how the effects of interruptions on working memory tasks can be mitigated by providing the opportunity to self-determine the time point of resumption, and how having further time to refocus attention before the resumption could help people recover after an interruption. Participants were asked to complete a visual working memory task that was interrupted randomly, after which they could either have additional time or decide on their own when to resume. By marking this time of resumption for the working memory task with a retrospective cuing paradigm, we either provided additional time or the chance to self-determine when to resume after interruptions and before the presentation of the retro-cue. In a blocked setting, participants were informed if they would be allowed to flexibly decide when to resume, or if they would be asked to resume on a fixed timing. We measured behavioural performance such as task accuracy and response times for both the primary working memory task and the interrupting task. Furthermore, we recorded the EEG and focused on neural oscillations as correlates of the underlying cognitive processes during the maintenance phases and resumption phases following the interrupting task. We argue that having more time to recover after an interruption or being able to pace the task resumption on your own should help minimising the deficits in primary task performance arising from interruptions.

As expected, interruptions overall reduced main task performance which could be seen as higher angular errors produced during the working memory task (see Figure 2A), and a general slowing of responses to this task (see Figure 2B). This is in line with the previous studies which showed decreased main task performance due to interruptions (Arnau et al., 2019; Clapp et al., 2010; Mishra et al., 2013; Rösner, Zickerick, et al., 2022; Ülkü et al., 2024; Zickerick, Kobald, et al., 2021). Further, even though working memory accuracy did not differ between different duration types, we were able to see an effect of longer resumption phases on response times, regardless of whether they were interrupted or not. Additionally, the flexible resumption condition provided a significant benefit over interruptions with fixed resumption phases in terms of working memory task response times, which made them on par with no-interruption conditions (see Figure 2B). Thus, we can say that in the case of self-determined resumption, participants were able to refocus their attention back to the main task on their own terms. Furthermore, when the main task accuracy was tested against the duration of this self-determined resumption, we observed that participants performed better when they took as little time as possible following an interruption. More simply, participants did not perform better by having more time after an interruption, in the case of the flexible condition. Instead, their performance showed a trend where it was better when they did not spend so much time to resume the main task, while still showing a general benefit of a self-determined resumption phase. When the main task response times were also tested the same way against the latency of the resumption, we saw that there was no clear relation between the two parameters, i.e. participants were not faster to respond by having more time to prepare their response. Hence, we were able to say that this self-determination benefit does not simply arise from taking their time and preparing as much as possible, because in both behavioural measures we don’t see an increase in performance by taking more time to resume. The flexibility itself enables participants to shift the focus of their attention more efficiently.

On an electrophysiological level, we were able to see that the interruptions resulted in weaker alpha/beta suppression and weakened evoked theta to the retro-cue compared to the same oscillatory measures observed in no-interruption conditions (see Figure 4). We interpret the observed effects as a deficit in the re-orienting of attention to the interrupted task, since previous studies showed increased theta power and stronger alpha/beta suppression when focusing attention on the level of working memory (De Vries et al., 2018; Rösner, Zickerick, et al., 2022; Schneider et al., 2017). It can thus be assumed that having to deal with an interrupting task during working memory storage exhausted available resources for an efficient focusing of attention on task-relevant information in working memory. Furthermore, we observed separate effects on alpha and beta suppression between shorter and longer duration trials (see Figure 4), with the latter showing stronger beta suppression before and around the onset of the retro-cue, and a sustained alpha power increase starting before the onset of the retro-cue and dissipating around retro-cue onset. Since we know that interruptions disturb the reactivation of motor representations in working memory (Ülkü et al., 2024; Zickerick, Kobald, et al., 2021), these effects might reflect preparatory mechanisms for the resumption of the working memory task. The longer duration trials, regardless of whether they contained an interruption or not, provided an overall advantage because participants were able to maintain the elements of the task robustly and prepare the yet-to-be executed response to the working memory task.

In addition to these main effects, we also observed an interaction between interruptions and the duration before resumption. Here, we saw three significant clusters with distinct topographies that arose from this interaction. Post-hoc testing revealed that the interaction within the oscillatory beta band was mainly driven by the interruption conditions, showing stronger beta suppression for longer resumption phases, with a distinct topography over the motor cortex contralateral to the response hand. We consider this as a marker of motor preparation for the working memory task response (Schneider et al., 2017; Van Ede et al., 2019; Zickerick, Kobald, et al., 2021), indicating that when given enough time, participants prepare better for the resumption of the main task. This point is further solidified by the relation between working memory task accuracy and oscillatory power in this cluster. Only performance after longer resumption phases benefited from stronger beta suppression. We argue that this effect is a compensatory mechanism to ease the interruption deficits on the working memory task performance. Hence, participants could not make use of this compensatory reactivation of motor planning during short interruption trials since they were not provided enough time for such mechanisms. For the oscillatory alpha band, on the other hand, we saw increased alpha power for longer resumption phase trials, which was even stronger for interruption trials with longer resumption phases, compared to the no-interruption condition with the same duration. This seems to be related to the well-known effect that alpha power increases during the maintenance phase of a working memory task when the task-relevant information needs to be protected from interference (Clayton et al., 2018; Johnson et al., 2011; Ülkü et al., 2024), potentially reflecting the active storage of relevant information in working memory. Here, such a mechanism might provide an additional compensatory mechanism for interruptions with longer resumption phases Additionally it reflects the refocusing of attention on task-relevant information in working memory even before the presentation of the retro-cue when more time is provided. We also see an effect on oscillatory theta band before the retro-cue, which shows reduced theta power for longer than for shorter resumption phases. When the general oscillatory patterns (see Figure 3) are considered, we can see that this effect arises from the theta oscillatory response induced by the interrupting task which decreases when participants realize that they have additional time to resume the main task (in the long resumption phase condition). We thus argue that this effect might reflect participants disengaging attention from the interruption task to reallocate available mental resources back to the main task.

Finally, we also observed a stronger alpha/beta suppression after the retro-cue for flexible resumption compared to fixed resumption phases. Post-hoc testing on this effect revealed that the alpha/beta suppression was significantly stronger for flexible resumption trials than for short and long resumption trials, which suggests that attentional focusing on the cued working memory content proceeded more efficiently. More interestingly, flexible resumption trials showed no difference when compared to the no-interruption conditions. Thus, we can say that being able to self-determine the resumption of the visual working memory task enabled participants to partly compensate for the deficits in orienting attention to task-relevant information after an interrupting task. This is not just about providing more time for resumption, but about the chance to disengage from the interrupting task and shift the focus of attention back to the main task in a self-determined way. This in turn could be seen in the working memory task response times that are comparable to the no-interruption conditions. Furthermore, when the oscillatory patterns were analysed as a function of working memory task accuracy, we could observe that participants benefitted from stronger alpha/beta suppression in the very same time-frequency cluster (see Figure 6). Although we found no direct difference in working memory performance between the flexible resumption condition and the conditions with fixed resumption phases, this may indicate that the self-determined resumption of a primary task after an interruption can facilitate attentional control processes, which in turn positively affect working memory performance.

Summarized, interruptions decrease performance in the interrupted task, but this issue can be mitigated by being able to self-determine when to resume the main task after having been interrupted. This particularly concerns the speed of responding in the primary task. Further, having additional time after an interruption can help people refocus their attention more efficiently, thereby mitigating deficits arising from interruptions. Finally, flexibility seems to provide additional benefits compared to just having more time for main task resumption after an interruption, as the oscillatory markers for attentional selection and action preparation benefit from self-determination, which in turn translates to a better working memory task accuracy as was seen in median-split data.

Even though we aimed for a more ecologically valid and real-world applicable paradigm, with the possibility to flexibly resume the main task after an interruption, the design and the tasks we used are still quite artificial and the findings might not directly be transferrable to application. However, the aim of this study was rather to gain an understanding of the underlying cognitive mechanisms and possible compensatory processes when processing interruptions. We are thus able to conclude that there are ways to cope with deficits arising from being interrupted. If interruptions cannot be avoided all-together, sufficient time or, ideally, temporal flexibility should be provided for primary task resumption.

## Data Availability Statement

Codes that are used for analysis are publicly available on the Open Science Framework (OSF) platform from the time of publication: https://osf.io/qw94r/?view_only=fc39302417424ae09894e8caecb0fe2c (a link to the data has to be requested).

## Author contributions

**S.Ü., S.G. and D.S.** designed the experiment. Data collection was performed by **S.Ü.** The data analysis and interpretation were done by **S.Ü.** under the supervision of **S.G., E.W. and D.S. S.Ü., S.G. and D.S.** drafted the manuscript, and all authors contributed to the revision of the manuscript and approved the final version of the manuscript for submission.

## Acknowledgments

We would like to thank Tobias Blanke for his support in programming the experiment, our entire laboratory staff for their support with the data recording, and two anonymous reviewers for their critical review and constructive comments on the manuscript. This work was supported by a third-party funding project from the German Research Foundation to D.S. (https://gepris.dfg.de/gepris/projekt/458108186).

